# Homologies and evolution of male tail characters in rhabditid and diplogastrid nematodes

**DOI:** 10.1101/2023.11.22.568293

**Authors:** Karin Kiontke, Simone Kolysh, Rocio Ng, David H. A. Fitch

## Abstract

A major question in evolutionary biology is how often the same developmental events, mechanisms and genes are reused in the recurrent evolution of similar phenotypes. If this happens frequently, it would suggest that evolution is often constrained by developmental genetic mechanisms. To help address this question, we used adherens junction staining and laser ablation to analyze the development underlying several features of nematode male tails have evolved recurrently. We find that recurrent evolution has sometimes employed similar developmental events (parallel evolution) and sometimes different events (convergent evolution). Specifically, phasmid position changed four times via cell migration and never by switches in cell lineage polarity; different genital papillae are missing in species with less than nine; and tail tip morphogenesis was gained at least twice (once with tail tip cell fusions and once without) and lost at least twice. As in previous analyses, we also find that genital papilla positions have shifted differently in different lineages relative to their conserved positions of origin in the lateral hypodermis. In particular, the v1 papilla homolog in diplogastrids has moved dorsally relative to the other v-papillae and lies posterior to the v2 papilla. The prevalence of recurrently evolved characters (homoplasy) suggests that caution should be exercised when using these characters for phylogenetic inference. On the other hand, because of their recurrent evolution, these characters provide good models for investigating how developmental and genetic systems may bias, constrain or allow phenotypic evolution.

## Results and Discussion

A major question in evolutionary biology is how often the same developmental events, mechanisms and genes are reused to make similar phenotypes (Stern & Orgogozo, 2008, 2009; Gompel & Prud’homme, 2009; Stern, 2011; Pereira & Kohlsdorf, 2023). That is, how strongly do developmental genetic mechanisms constrain or bias the evolutionary pathways leading to adaptive morphologies? To address this question, we can study the developmental events underlying phenotypes that have evolved recurrently in independent lineages. Here, we show that several male tail characters in rhabditid nematodes have evolved repeatedly to make similar phenotypes. We then tested if similar developmental events have been co-opted (parallelism), indicating possible constraint, or if the similar phenotypes result from different developmental events (convergence), indicating plasticity. We found cases for both types of recurrent evolution.

### Recurrently evolving characters

The first step was to determine how many evolutionary changes in these characters occurred and in what species lineages. We chose species representing several different phylogenetic clades of rhabditid nematodes, plus a diplogastrid and two outgroup representatives (Fig. 1). Some of these species had been previously studied, i.e. the rhabditids *Metarhabditis blumi, Oscheius myriophilus, Reiterina typica, Rhabditella axei, Caenorhabditis elegans, Teratorhabditis palmara*, and *Pelodera strongyloides*, and outgroup representative *Panagrellus redivivus* (Fitch & Emmons, 1995; Fitch, 1997; Kiontke & Sudhaus, 2000; Sternberg & Horvitz, 1982). Here, we additionally studied rhabditids *Cruznema tripartitum, Rhabditoides inermis, Haematozoon subulatum*, and *Poikilolaimus oxycercus*, diplogastrid *Diplogasteroides nasuensis*, and outgroup representative *Brevibucca saprophaga*. Parsimony was used to trace character evolution on a robust cladogram (Fig. 1A) previously inferred from molecular data (Kiontke et al., 2007). Recurrent evolution was found for the following characters: the phasmid switched at least three times from a posterior to a more anterior position, a genital papilla (GP, ray) was lost at least twice, and tail tip morphogenesis (TTM) was gained at least twice (in concert with the appearance of a bursa or fan) and lost at least twice (yet retaining at least a narrow fan). We focused our developmental analyses on the latter three characters.

**Figure 1.**
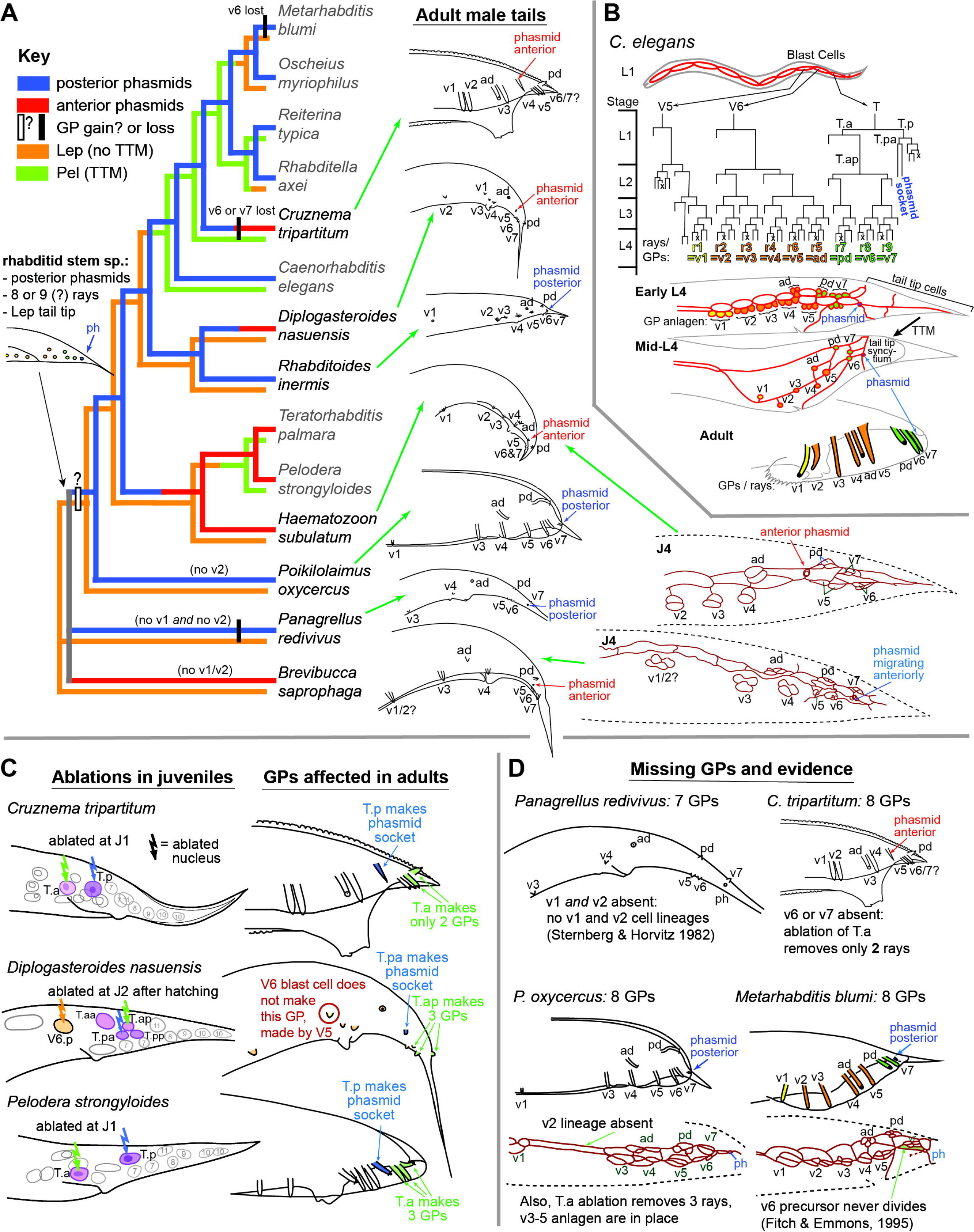
Evolution of male tail characters in rhabditid nematodes. (**A**) Cladogram of representative species or Rhabditidae plus two outgroup representatives (*Panagrellus redivivus* and *Brevibucca saprophaga*), based on molecular analyses (Kiontke et al., 2007). Left side views of adult male tails (redrawn based on Kiontek & Sudhaus, 2000) are shown for the species investigated here (species names in black type); species with names in gray have been investigated previously (Fitch & Emmons, 1995; Kiontke & Sudhaus, 2000; with names revised as per Sudhaus, 2011). Evolution of three characters is traced by parsimony on the cladogram with character states noted in the key: phasmids posterior (blue lineages) versus anterior (red lineages), tail tip peloderan (Pel, green lineages) versus leptoderan (Lep, orange lineages), and gain (open rectangle) versus loss (filled rectangle) of a ray. Question marks and gray lineages represent ambiguous assignment of character states or changes. The inferred rhabditid stem species’ male tail likely had posterior phasmids, no fan (bursa), and a Lep tail tip, but it is ambiguous whether it had 8 or 9 genital papillae (GPs, rays) per side. Early J4 adherens junction staining patterns are shown for *Haematozoon subulatum* (showing the phasmid already in the “anterior” position by this stage) and *Brevibucca saprophaga* (showing the phasmid beginning to move anteriorly relative to the posterior GPs only at this stage). (**B**) The *C. elegans* cell lineages of the three blast cells (V5, V6 and T) that make the rays (GPs) and phasmids. Three cells make each ray: one becomes the “structural cell” and the other two become neurons (). By convention, juvenile stages are designated L1–L4 in *C. elegans*, but J1–J4 in other species. After the anlage for the GPs originate in the lateral hypodermis in L3, the ray cells cluster (early L4), a conserved pattern that allows assignment of ray homologies across species (Fitch & Emmons, 1995). Rays then differentiate, leaving the ray structural cells at the lateral surface (mid-L4) at about the same time as tail tip morphogenesis occurs (Nguyen et al., 1999). After male tail morphogenesis is complete, the adult has finger-like rays in the same anteroposterior order as the structural cells in the mid-L4 stage (Fitch & Emmons, 1995). (**C**) Depicted are the cells that were ablated in J1 (*Cruznema tripartitum* and *Pelodera strongyloides*) or J2 (*Diplogasteroides nasuensis*) stages (left) and the GPs in adults that were affected by the ablations (right). See text for details. (**D**) Evidence for assigning homologies to the “missing GPs” of four species: *Panagrellus redivivus* (v1 and v2 cell lineages are absent; Sternberg & Horvitz, 1982), *C. tripartitum* (ablation of T.a only removes 2 rays, one of which is ‘pd’, suggesting that v6 or v7 is absent), *Poikilolaimus oxycercus* (v2 cell lineage is absent, v3-5 anlagen are in place and T.a ablation removes 3 posterior rays, all consistent with v2 being absent), and *Metarhabditis blumi* (the v6 ray is missing because the v6 precursor fails to divide; Fitch & Emmons, 1995).

### Phasmid position

Using dyes selectively taken up by the chemoreceptive phasmids but not the mechanosensory rays (genital papillae, GPs) and scanning electron microscopy, we showed previously that phasmids have changed position relative to GPs (Fitch & Emmons, 1995; Kiontke & Sudhaus, 2000). Specifically, whereas the phasmid is posterior of all GPs in some species (e.g. rhabditids *C. elegans, M. blumi, O. myriophilus, R. typica, R. axei*, and *P. oxycercus*, and outgroup taxon *P. redivivus*, it is anterior of the most-posterior GPs in other species (e.g. rhabditids *R. inermis, C. tripartitum, T. palmara, P. strongyloides*, and *H. subulatum* (Fitch & Emmons, 1995; Kiontke & Sudhaus, 2000). For shorthand, we call these “posterior” versus “anterior” phasmid positions. Here, we confirmed that the diplogastrid *D. nasuensis* has anterior phasmids and discovered that outgroup taxon *Brevibucca saprophaga* does too.

Tracing the evolution of this character on the cladogram shows that anterior phasmids arose three times independently in rhabditids from ancestors having posterior phasmids: in the lineages to *Cruznema*, diplogastrids and the clade containing *T. palmara* and *P. strongyloides* (Pleiorhabditis) plus *H. subulatum* (Fig. 1A). *Brevibucca* also has anterior phasmids independently of the rhabditids, but whether this was the primitive (plesiomorphic) or derived (apomorphic) state cannot be inferred from this cladogram.

In *C. elegans*, the bilateral T blast cells (TL and TR) each produce a phasmid socket (from posterior daughter cell T.p) and the posterior three GPs (from anterior daughter cell T.a) (Fig. 1B). In *lin-44/Wnt* mutations, this polarity is reversed, such that phasmid socket-like lineages are produced from T.a and GP-like lineages from T.p (Herman & Horvitz, 1994). Thus, one hypothesis for a developmental event switching phasmid position from posterior to anterior is a polarity reversal of the first T cell division, such as might be produced by a heterochronic delay in the expression of the LIN-44/Wnt signaling ligand (or its receptor) relative to its expression in *C. elegans* (Fitch 1997; Kiontke & Sudhaus 2000). An alternative hypothesis is that no such reversal occurs and instead the cells producing the phasmid socket migrate anteriorly from an originally posterior position.

To discriminate between these hypotheses, we laser-ablated either T.a or T.p in J1 juveniles of *C. tripartitum* and *P. strongyloides* to test if this affected either the appearance of the phasmid or the GPs in adults (Fig. 1C). Additionally, we ablated T.ap or T.pa in J2 juveniles of *Diplogasteroides nasuensis*; in this species, the J1 molt occurs inside the eggshell and J2s subsequently hatch, making ablations difficult in J1s. In all cases, ablating T.p (or T.pa) resulted in the deletion of only the phasmids and ablating T.a (or T.ap) deleted only GPs. Thus, all three instances of recurrent evolution of anterior phasmids in rhabditids involve cell migration. Additionally, by following the origins of GP cells by staining adherens junctions, we found that all GP cell lineages occur anterior to the phasmid in *Brevibucca saprophaga*, but the phasmid socket migrates anteriorly during J4 development. In the rhabditids, the MH27 staining patterns show that the phasmid socket cells are already at the anterior position prior to J3.

These results suggest that there is a constraint on how anterior phasmids can form. For example, variation in a signaling pathway like Wnt that affects T blast-cell polarity, but which is also used during embryogenesis and in other developmental fate decisions, may tend to produce pleiotropic consequences and thus would tend to be selected against. However, the timing of phasmid migration appears to be plastic.

### Genital papillae

By staining the adherens junctions surrounding epidermal cells that produce the GPs and phasmid sockets, we showed previously that GPs originate from anlagen arising in a characteristic, archetypal pattern that is conserved in rhabditids (Fig. 1B; Fitch & Emmons 1995; reviewed in Haag et al. 2018). Because of this conservation, homologies can be assigned to each GP across different species, despite species-specific shifts in GP positions after their origin (Fitch & Emmons 1995; Fitch 1997). A naming system for GP homologs was adopted (Sudhaus & Fürst von Lieven 2003) whereby the seven ventrolateral GPs on each side are designated “v1–v7”, counting anterior to posterior, and the two subdorsal GPs are designated “ad” and “pd” (for “anterior dorsal” and “posterior dorsal”). Along the anteroposterior axis, the order of GP anlagen is v1,v2,v3/v4,ad,v5,v6,v7,pd,ph (“ph” = phasmid; “/” indicates position of the cloaca). Although the cells forming the “v” GPs tend not to shift ordinal position relative to each other, they do migrate to different positions ventral of the subdorsal GPs ad and pd, resulting in different anteroposterior orders of GPs in different species.

Using the same method to stain adherens junctions, we analyzed GP development in *D. nasuensis, H. subulatum, P. oxycercus*, and *B. saprophaga*. Fig. 1A shows the resulting homology assignments of GPs in adults. As an example, the homology formula for the GP/ray pattern in adult males of *H. subulatum* is v1,v2,v3/(v4,ad)/ph,(v5,pd),(v6,v7), where parentheses designate GP clusters and the two slashes in this case mean that v4 and ad are at the same position as the cloaca.

Interestingly, *D. nasuensis*—like other diplogastrids—has three dorsal papillae, not only two as in rhabditids. Specifically, in diplogasterids, a third dorsal GP lies anterior to the “ad” and “pd” GPs, and tends to lie in the second or third position along the anteroposterior axis. Its homology has remained an open question (Sudhaus & Fürst von Lieven 2000). Because we could not definitively answer this question with adherens junction staining, we laser-ablated the GP blast cells. In *C. elegans*, the V6 blast cell generates GPs v2–v5 and ad, the V5 blast cell generates v1, and the T blast cell generates pd, v6 and v7 in addition to the phasmid socket (Fig. 1B). We ablated V6 in *D. nasuensis* and found that only the anterior-most dorsal GP remained in the adult, which we infer must be v1. Thus, the GP homology formula for *D. nasuensis* is v2,v1/(v3,v4)/ad,ph,(v5,v6,v7,pd), where double slashes denote that GPs v3 and v4 are at the same anteroposterior position as the cloaca. Thus, in diplogasterids, v1 migrates dorsally, and other “v” GPs can then slide to a position anterior of v1. Interestingly, ray 1 (v1) in *C. elegans* also migrates a little dorsally and opens on the dorsal surface of the fan. It is possible that this GP has a tendency toward dorsal migration, realized in a more extreme way in diplogastrids.

Establishing GP homologies across these species allowed us to identify which GPs are missing in species with less than nine GPs (Fig. 1D). According to the cladogram, one bilateral pair of rays was lost in the lineages to 8-rayed *M. blumi* and *C. tripartitum. P. oxycercus* also has only 8 rays, as does outgroup representative *B. saprophaga*. The other outgroup representative, *P. redivivus*, has only 7 rays. It is therefore possible that the rhabditid stem species had 9 GPs and one was lost in the lineage to *Poikilolaimus*, or it had 8 GPs and one was gained in the ancestor to the rest of the rhabditids after divergence of *Poikilolaimus* (Fig. 1A). It was previously found that the R8 ray neuroblast fails to divide in *M. blumi* (Fitch & Emmons, 1995), identifying its missing GP as v6. In *C. tripartitum*, ablation of the T.a blast cell only removes the two posterior-most rays (pd and another ray), suggesting that either v6 or v7 was lost in this species lineage. Ablation of T.a in *P. oxycercus* results in removal of three posterior rays, so v6, v7 and pd homologs are present in this species. The adherens junction patterns also show that anlagen for v3–v5 are in their normal places; however, a large undivided cell intervenes between the v3 cell clusters and those of the most anterior GP, suggesting that v2 is the absent GP. Also, *P. redivivus* has only 7 rays; from the published cell lineage (Sternberg & Horvitz 1982), it is clear that both v1 and v2 are absent. Thus, in most cases, different rays are missing in different species. Because v2 is the missing ray in *P. oxycercus*, and v1 or v2 (or both) is missing in the outgroup species, the rhabditid stem species may not have had v2 homolog.

These results suggest that there is not strong constraint on which GPs can be lost. Genetic and ablation studies have demonstrated considerable redundancy among rays in *C. elegans* for aspects of male mating behavior; as few as 3 pairs of rays are required to allow successful mating, although robustness is enhanced with additional rays (Liu & Sternberg, 1995; Sherlekar & Lints, 2014). Thus, it is conceivable that loss of any particular ray pair may not matter much to fitness, although there have not been such rigorous study in gonochoristic species, which are expected to have higher levels of sperm competition.

### Tail tip morphogenesis

In some rhabditid species, male-specific tail tip morphogenesis (TTM, tail tip “retraction”) occurs during J4 to change the pointed tails of juveniles into rounded, “peloderan” (Pel) tails, whereas TTM does not occur in other species, leaving tail tips pointed and “leptoderan” (Lep), e.g. with the point protruding beyond the fan. Females in nearly all species retain the pointed shape of the juvenile. In *C. elegans*, this sexual dimorphism is governed by a gene regulatory network with DMD-3 at its hub (Mason et al. 2008; Nelson et al. 2011). The ancestral, plesiomorphic state for rhabditids is leptoderan; peloderan tails evolved independently at least twice and reverted to leptoderan tails at least twice independently (Fig. 1A; Kiontke & Fitch, 2005).

Previous transmission electron microscopy and analyses of adherens junctions showed that the tail tip cells hyp8–11 fuse during TTM in *C. elegans* (Nguyen et al. 1999), but not in the peloderan *R. typica, T. palmara* or *P. strongyloides* (Fitch 2000). Tail tip cells also do not fuse in leptoderan species *O. myriophilus, M. blumi*, and *R. axei*.

Here, we also found that tail tip cells remain unfused in leptoderan *D. nasuensis, R. inermis, H. subulatum, P. oxycercus, P. redivivus*, and *B. saprophaga*. Thus leptoderan tails seem to be made in similar ways, but cell fusions accompany tail tip morphogenesis only in *C. elegans*. Our data are still unclear as to whether or not tail tip cells fuse in the peloderan *C. tripartitum*.

### Summary

Recurrent evolution has occurred with respect to several male tail characters in rhabditid nematodes. Because of this “homoplasy”, these characters may not be useful for inferring species relationships. On the other hand, these recurrently evolving characters are interesting models for investigating the degree to which evolution may be constrained by developmental-genetic mechanisms. For example, change in phasmid position appears to be constrained, as it is accomplished only by cell migration and never by switches in cell polarity (cf. previous hypotheses in Fitch, 1997; Kiontke & Sudhaus, 2000), although the timing of this migration is plastic. However, evolution of other male tail characters is less constrained. For example, although the GP anlagen are produced in a highly conserved spatial pattern during J3–J4 development, developing GPs move to different positions in different species. In diplogastrids, v1 becomes an anterior, dorsal GP. Also, different GPs can be lost when the number of GPs is reduced below nine. Finally, male-specific tail tip morphogenesis may or may not involve cell fusions. Thus, nematode male tail morphogenesis offers multiple opportunities to investigate how gene regulatory networks have been conserved, re-used or diversified in the course of evolution. We are currently addressing this question for the recurrent evolution of TTM.

## Methods

### Animal husbandry

Strains of all species were kept on Nematode Growth Medium agar plates as described previously (Sulston & Hodgkin, 1988). The plates were seeded with a lawn of *E. coli* OP50-1, but other bacteria, possibly from the source habitats of the species were also present.

### Phylogenetic analysis

Using the cladogram published earlier for rhabiditds (Kiontke et al. 2007), we traced character evolution using the principle of parsimony, checking our inferences with Mesquite (Maddison & Maddison, 2023).

### Adherens junction immunofluorescence

As described previously (Fitch & Emmons, 1995), MH27 mouse monoclonal antibody (from supernatent of cells provided by R. Waterston) was used with rhodamine-conjugated goat anti-mouse secondary antibody (Sigma-Aldrich) to stain fixed animals of different stages.

### Laser ablation

Laser ablation of blast cells in the male tail was performed as described previously (Walston & Hardin, 2010) using a Zeiss Axioskop I with a 337-nm nitrogen pumping laser (Micropoint, Photonic Instruments) connected via optical cable to a dye cell module (at the epiillumination port) filled with coumarin 440 dye (5 mM in methanol). Targeting was performed using a video camera connected to a TV monitor. Ablated cells and the resulting adult phenotype were documented by drawings using a camera lucida attached to the microscope.

## Reagents

| Strain and Species Binomen | Genotype | Available from |
| --- | --- | --- |
| **DF5010 *Metarhabditis blumi*** | wild type | CGC (*Caenorhabditis* Genetics Ctr) |
| EM435 *Oscheius myriophilus* | wild type | CGC |
| DF5025 *Reiterina typica* | wild type | CGC |
| DF5006 *Rhabditella axei* | wild type | CGC |
| SB202 *Cruznema tripartitum* | wild type | NYURC (NYU Rhabditid Collection), Fitch lab |
| CB4088 *Caenorhabditis elegans* | *him-5(e1490) V* | CGC |
| SB335 *Diplogasteroides nasuensis* | wild type | NYURC |
| SB328 *Rhabditoides inermis* | wild type | NYURC |
| DF5019 *Teratorhabditis palmarum* | wild type | CGC |
| DF5022 *Pelodera strongyloides* | wild type | CGC |
| SB303 *Haematozoon subulatum* | wild type | NYURC |
| SB200 *Poikilolaimus oxycercus* | wild type | NYURC |
| PS1163 *Panagrellus redivivus* | wild type | CGC |
| SB261 *Brevibucca saprophaga* | wild type | NYURC |

## Acknowledgements

Many thanks to Walter Sudhaus for originally providing strains of most of the species used here and to the CGC (*Caenorhabditis* Genetics Center) for archiving strains. The CGC is funded by NIH Office of Research Infrastructure Programs (P40 OD010440).

## Author Contributions

KK performed laser ablations, helped to interpret MH27 staining results, supervised collection of data, provided illustrations, and edited the manuscript.

SK and RN (undergraduate students at New York University during the course of this work) performed MH27 staining and interpreted those results.

DF directed the project, supervised the undergraduates, obtained funding, and wrote the manuscript.

